# Incomplete influenza A virus genomes are abundant but readily complemented during spatially structured viral spread

**DOI:** 10.1101/529065

**Authors:** Nathan T. Jacobs, Nina O. Onuoha, Alice Antia, Rustom Antia, John Steel, Anice C. Lowen

## Abstract

Viral genomes comprising multiple distinct RNA segments can undergo genetic exchange through reassortment, a process that facilitates viral evolution and can have major epidemiological consequences. Segmentation also allows the replication of incomplete viral genomes (IVGs), however, and evidence suggests that IVGs occur frequently for influenza A viruses. Here we quantified the frequency of IVGs using a novel single cell assay and then examined their implications for viral fitness. We found that each segment of influenza A/Panama/2007/99 (H3N2) virus has only a 58% probability of being present in a cell infected with a single virion. These observed frequencies accurately account for the abundant reassortment seen in co-infection, and suggest that an average of 3.7 particles are required for replication of a full viral genome in a cell. This dependence on multiple infection is predicted to decrease infectivity and to slow viral propagation in a well-mixed system. Importantly, however, modeling of spatially structured viral growth predicted that the need for complementation is met more readily when secondary spread occurs locally. This expectation was supported by experimental infections in which the level spatial structure was manipulated. Furthermore, a virus engineered to be entirely dependent on co-infection to replicate *in vivo* was found to grow robustly in guinea pigs, suggesting that coinfection is sufficiently common *in vivo* to support propagation of IVGs. The infectivity of this mutant virus was, however, reduced 815-fold relative wild-type and the mutant virus did not transmit to contacts. Thus, while incomplete genomes augment reassortment and contribute to within-host spread, the existence of rare complete IAV genomes may be critical for transmission to new hosts.

---

Pathogen evolution poses a continued threat to public health by reducing the effectiveness of antimicrobial drugs and adaptive immunity. In the case of the influenza A virus (IAV), this evolution results in seasonal outbreaks as new viruses emerge to which pre-existing immunity is weak. Each year requires a new vaccine as a consequence, and keeping pace with IAV evolution is challenging: unexpected emergence of new strains could render the vaccine obsolete before the flu season starts. IAV populations evolve rapidly in part because their mutation rates are high, on the order of 10^−4^ substitutions per nucleotide per genome copied^1^. The segmentation of the viral genome gives a second source of genetic diversity. The IAV genome is composed of eight single-stranded RNA segments, and so cells co-infected with two different IAV virions can produce chimeric progeny with a mix of segments from these two viruses. This process, termed reassortment, carries costs and benefits analogous to those of sexual reproduction in eukaryotes^2^. Reassortment can join beneficial mutations from different backgrounds to alleviate clonal interference^3^, and purge deleterious mutations to mitigate the effects of Muller’s ratchet^4, 5^. This combinatorial shuffling of mutations may accelerate adaptation to new environments such as a novel host^6^. But free mixing of genes through reassortment may also reduce viral fitness by separating beneficial segment pairings, as sexual reproduction carries this cost in eukaryotes^7^. Previous work has shown that reassortment occurs readily between closely related variants^8^, but is limited between divergent lineages due to molecular barriers^9, 10^ or reduced fitness of progeny^11, 12^. Nevertheless, the contribution of reassortment to emergence of novel epidemic and pandemic IAVs has been documented repeatedly^13-16^. Factors that affect the frequency of co-infection and consequently reassortment are therefore likely to play an important role in viral evolution.

While the ability of a virus particle to enter a cell depends only on the proteins that line the virion surface, subsequent production of viral progeny requires successful expression and replication of the genome. A virion that does not contain, or fails to deliver, a complete genome could therefore infect a cell but fail to produce progeny. IAV particles outnumber plaque-forming units (PFUs) by approximately 10–100 fold^17^, meaning that only a minority of particles establish productive infection at limiting dilution. Recent data suggest that IAV infection is not a binary state, however. Efforts to detect viral proteins and mRNAs at the single cell level have revealed significant heterogeneity in viral gene expression^18–20^. These data furthermore suggest that a subset of gene segments is often missing entirely from cells infected at low MOI. Thus, many non-plaque-forming particles appear to be semi-infectious, giving rise to incomplete viral genomes (IVGs) within the infected cell^21^.

Replication and expression of only a subset of the genome may be explained by two potential mechanisms: either the majority of particles lack one or more genome segments, or segments are readily lost in the process of infection before they can be replicated. Electron microscopy has shown that most particles contain eight distinct RNA segments^22^, and FiSH-based detection of viral RNAs indicated that a virion tends to contain one copy of each segment^23^, suggesting that most particles contain full genomes. Regardless of the molecular mechanisms that lead to the phenomenon of incomplete IAV genomes, their frequent occurrence suggests that complementation by co-infection at the cellular level is an underappreciated aspect of the viral life cycle. The observation of appreciable levels of reassortment following co-infection at low MOIs suggests IVG reactivation through complementation occurs commonly during IAV infection^24^. Nevertheless, the extent to which IAVs rely on co-infection for replication, and how this need changes over the course of infection, remains unclear. Similarly, the existence of IVGs *in vivo* has been demonstrated^25^, but their importance to the dynamics of infection within hosts is untested.

Here we investigate the biological implications of incomplete IAV genomes and the emergent need for cooperation at the cellular level. We first developed a novel single-cell sorting assay to measure the probability of each segment being delivered by an individual virion for influenza A/Panama/2007/99 (H3N2) [Pan/99] virus. Our data estimate that individual virus particles lead to successful replication of all eight gene segments only 1.3% of the time. When considering a well-mixed system in which virus particles are distributed randomly over cells, the potential fitness costs of incomplete genomes are high. In contrast, a model of viral spread that incorporates local dispersal of virions to nearby cells predicted that the spatial structure of virus growth mitigates costs of genome incompleteness. Testing of this model confirmed that infections initiated with randomly distributed inocula contained more IVGs than those generated by secondary spread from low MOIs, in which spatial structure is inherent. To determine the potential for complementation to occur *in vivo*, we generated a mutant virus that was fully dependent on cellular co-infection for viral replication, and found that it was able to grow within guinea pigs, but unable to transmit to cagemates. Taken together, these results suggest that the abundance of incomplete genomes and the potential for complementation are important factors in the replication and transmission of IAV.

## Results

### Measurement of P_P_

To better evaluate the implications of genome incompleteness for IAV fitness and reassortment, we sought to quantify the probability of successful replication (P_P_ = probability present) for each of the eight IAV genome segments within single cells infected with single virus particles. To ensure accurate detection of IVGs, we devised a system that would allow their replication to high copy number. We applied our approach to the human seasonal isolate, influenza A/Panama/2007/99 (H3N2) virus. In this assay, MDCK cells are inoculated with a virus of interest, referred to herein as “Pan/99-WT” or “WT”, and a genetically tagged helper virus (“Pan/99-Helper” or “Helper”). This Helper virus differs from the WT strain only by silent mutations on each segment that provide distinct primer-binding sites. For example, qPCR primers targeting WT PB2 will not anneal to cDNA of Helper PB2, and vice versa. By co-inoculating cells with a low MOI of WT virus and a high MOI of Helper virus, we ensure that each cell is productively infected, but is unlikely to receive more than one WT virion. Following infection, one cell per well is sorted into a 96-well plate containing MDCK cell monolayers. The initially infected cell produces progeny which then infect neighboring cells, effectively amplifying the vRNA segments present in the first cell. The presence or absence of WT segments in each well can then be measured by performing segment-specific RT qPCR. As detailed in the Methods, a correction factor was applied to account for multiple infection, the probability of which could be estimated based on the observed number of cells infected with each virus.

Using this assay, the P_P_ values for each segment of Pan/99 virus were quantified (Fig. 1A). We observed that each segment was present at an intermediate frequency between 0.5 and 0.6, indicating that incomplete genomes may arise from loss of any segment(s). The mean frequency across all segments was 0.58. When used to parameterize a model that estimates the frequency of reassortment, which we published previously^24^, these P_P_ values generated predicted levels of reassortment that align closely with experimental data (Fig. 1B). This match between observed and predicted reassortment is important because i) it offers a validation of the measured P_P_ values and ii) it indicates that IVGs fully account for the levels of reassortment observed, which are much higher than predicted for viruses with only complete genomes^24^.

**Figure 1.**
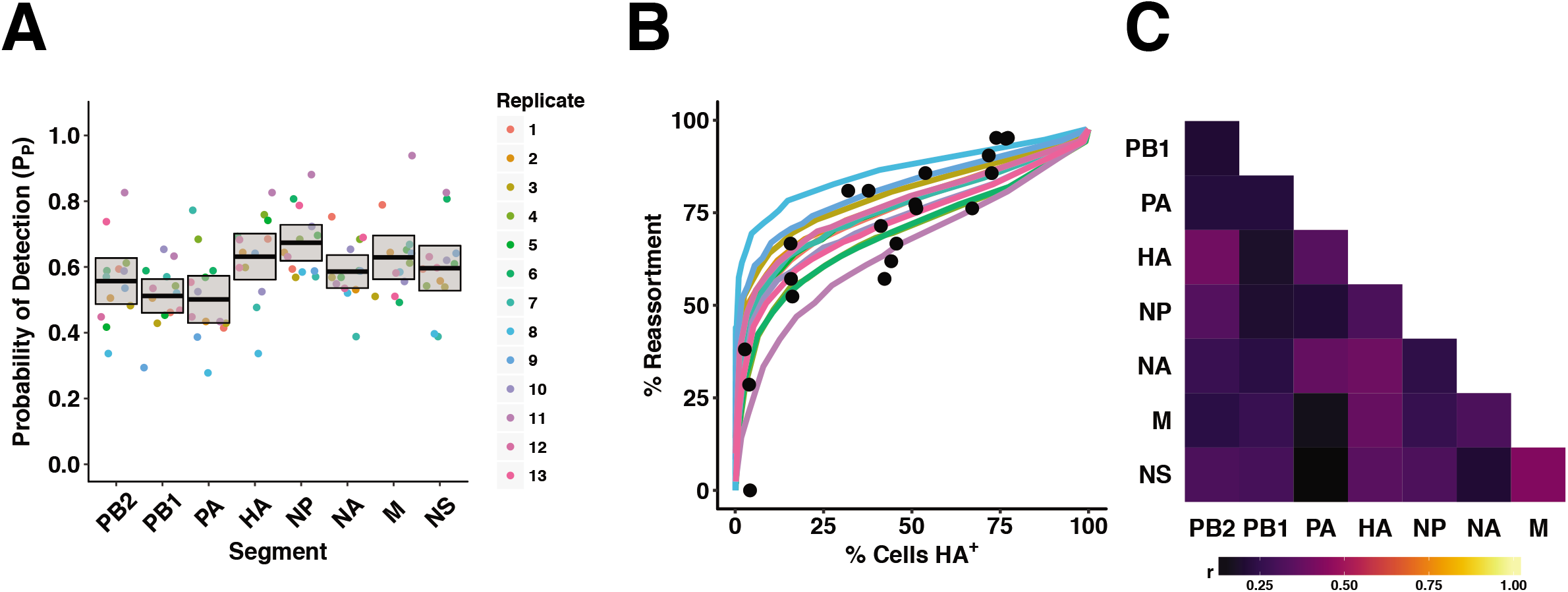
Incomplete genomes are common in Pan/99 virus infection. (A) Segment-specific P_P_ values were measured by a single-cell sorting assay. Each set of colored points corresponds to eight P_P_ values measured in a single experimental replicate, with thirteen independent replicates performed. Horizontal bars indicate the mean and shading shows the 95% CI. Segments encoding the polymerase subunits PB2, PB1, and PA were present significantly less frequently than others (p < 0.001, ANOVA). (B) Using each replicate’s P_P_ values as input parameters, the computational model from Fonville et al. was used to predict the frequency of reassortment across multiple levels of infection^24^. Black circles represent experimental data from Fonville et al. and show levels of reassortment observed following single cycle coinfection of MDCK cells with Pan/99-WT and a Pan/99 variant viruses. Colors correspond to the legend shown in panel A. (C) Pairwise correlations between segments (r) are shown as color intensities represented by a color gradient (below).

Interactions between vRNP segments are thought to play an important role in the assembly of new virions^10, 26–29^. To determine whether similar interactions exist that could mediate the co-delivery of segments to the cell, the patterns of segment co-occurrence were analyzed. In performing this analysis, it was again important to take into account the known probability of multiple infection in our single cell assay. As shown in Supplementary Figure 1, cells containing more segments were likely to have been infected with multiple virions. Because such cells are less informative for this analysis, we applied a weighting factor to ensure that results relied more strongly on data from cells with fewer WT segments. Namely, we determined the probability that a given cell acquired its gene constellation by infection with a single virion and weighted data according to this probability to calculate the pairwise correlation between segments. While some significant interactions were observed (HA-NA, HA-M, M-NS), they were relatively weak, with r^2^ values below 0.15 (Fig.1C). Thus, our data suggest that associations among specific vRNPs do not play a major role during the establishment of infection within a cell.

### Predicted costs of incomplete genomes for cellular infectivity

If singular infections often result in replication of fewer than eight viral gene segments, the infectious unit would be expected to comprise multiple particles. To evaluate the relationship between the frequency of IVGs and the number of particles required to infect a cell, we developed a probabilistic model in which the likelihood of segment delivery is governed by the parameter P_P_. In Figure 2A we examine how P_P_ affects the frequency with which a single virion delivers a given number of segments. If P_P_ is low, singular infections typically yield few segments per cell. Even at the intermediate P_P_ that characterizes Pan/99 virus, the vast majority of singular infections give rise to IVGs within the cell. When P_P_ is high, however, most cells receive the full complement of eight segments. In Figure 2B we plot the relationship between P_P_ and the percentage of cells that are expected to be productively infected following singular infection. If only a single virus infects a cell, then the probability that all eight segments are present will be P_p_^8^. For Pan/99 virus, the frequency with which eight segments are present is approximately 0.58^8^ = 1.3%.

**Figure 2.**
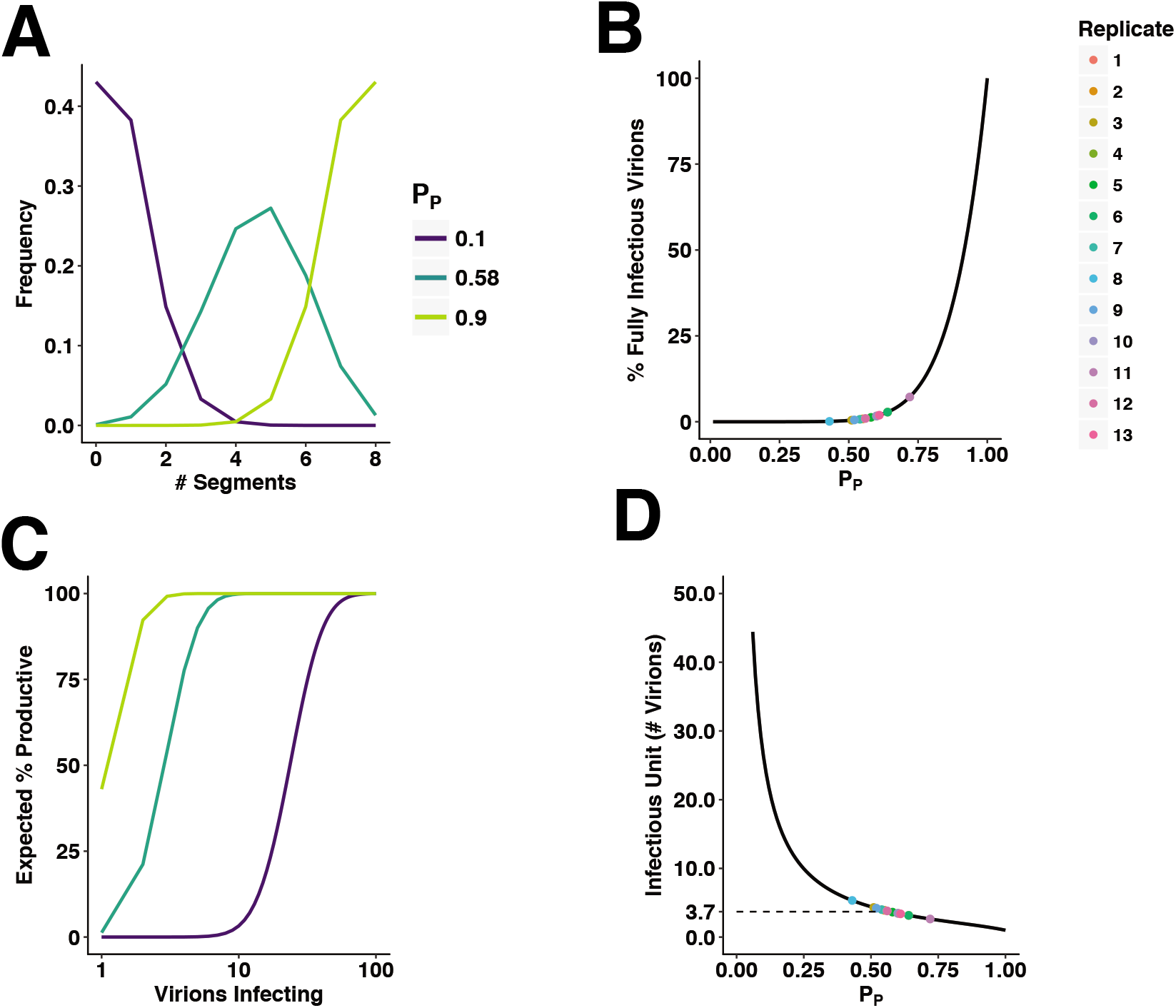
Incomplete genomes require complementation for productive infection at the cellular level. (A) The expected number of segments delivered upon infection with a single virion was calculated for two extreme values of P_P_ (0.10, 0.90) and the estimated P_P_ of Pan/99 virus (0.58). (B) The percentage of virions expected to initiate productive infection was plotted as a function of P_P_. Colored points correspond to the average P_P_ value of each experimental replicate in Fig. 1 and therefore show the prediction for Pan/99 virus. (C) The probability that a cell will be productively infected following infection with a given number of virions was calculated for the same P_P_ values as in (A). (D) The expected number of virions required to make a cell productively infected is plotted as a function of P_P_. As in (B), colored points correspond to the average P_P_ value of each Pan/99 experimental replicate in Fig. 1.

Importantly, however, if more than one virus particle infects the cell, then the probability that all eight segments are present will be considerably higher. This effect is demonstrated in Figure 1C, where the percentage of cells containing all eight IAV segments is plotted as a function of the number of virions that have entered the cell. Here we see that, even for low P_P_, a high probability of productive infection is reached at high multiplicities of infection. Finally, in Figure 2D, the relationship between P_P_ and the average number of virions required to productively infect a cell is examined. We see that the number of virions comprising an infectious unit is inversely proportional to P_P_. Based on our experimentally determined values of P_P_ for Pan/99 virus, we estimate that an average of 3.7 virions must enter a cell to render it productively infected (Fig. 2D). Thus, as a result of stochastic loss of gene segments, the likelihood that a full viral genome will be replicated within a singularly infected cell is low. The fitness implications of this inefficiency may be offset, however, by complementation of IVGs in multiply infected cells.

### Predicted costs of incomplete genomes for population infectivity

The potential for multiple infection to mitigate the costs of inefficient genome delivery will, of course, depend on the frequency of multiple infection. To evaluate the theoretical impact of IVGs on viral fitness, we therefore modeled the process of infection at a population level. A population of computational virions was randomly distributed across a population of computational cells over a range of MOIs, such that the likelihood of multiple infection was dictated by Poisson statistics. The frequency with which each cell acquired segments was again governed by P_P_. For each combination of P_P_ and MOI, we calculated the percentage of populations in which at least one cell contained eight segments (Fig. 3A). This plot shows that viruses with lower P_P_ require markedly higher MOIs to ensure productive infection within a population of cells. Indeed, when we estimated the MOI required for a virus of a given P_P_ to infect 50% of populations, we observed that the ID_50_ increases exponentially as P_P_ decreases (Fig. 3B). Thus, a reliance on multiple infection in a well-mixed system is predicted to bear a substantial fitness cost.

**Figure 3.**
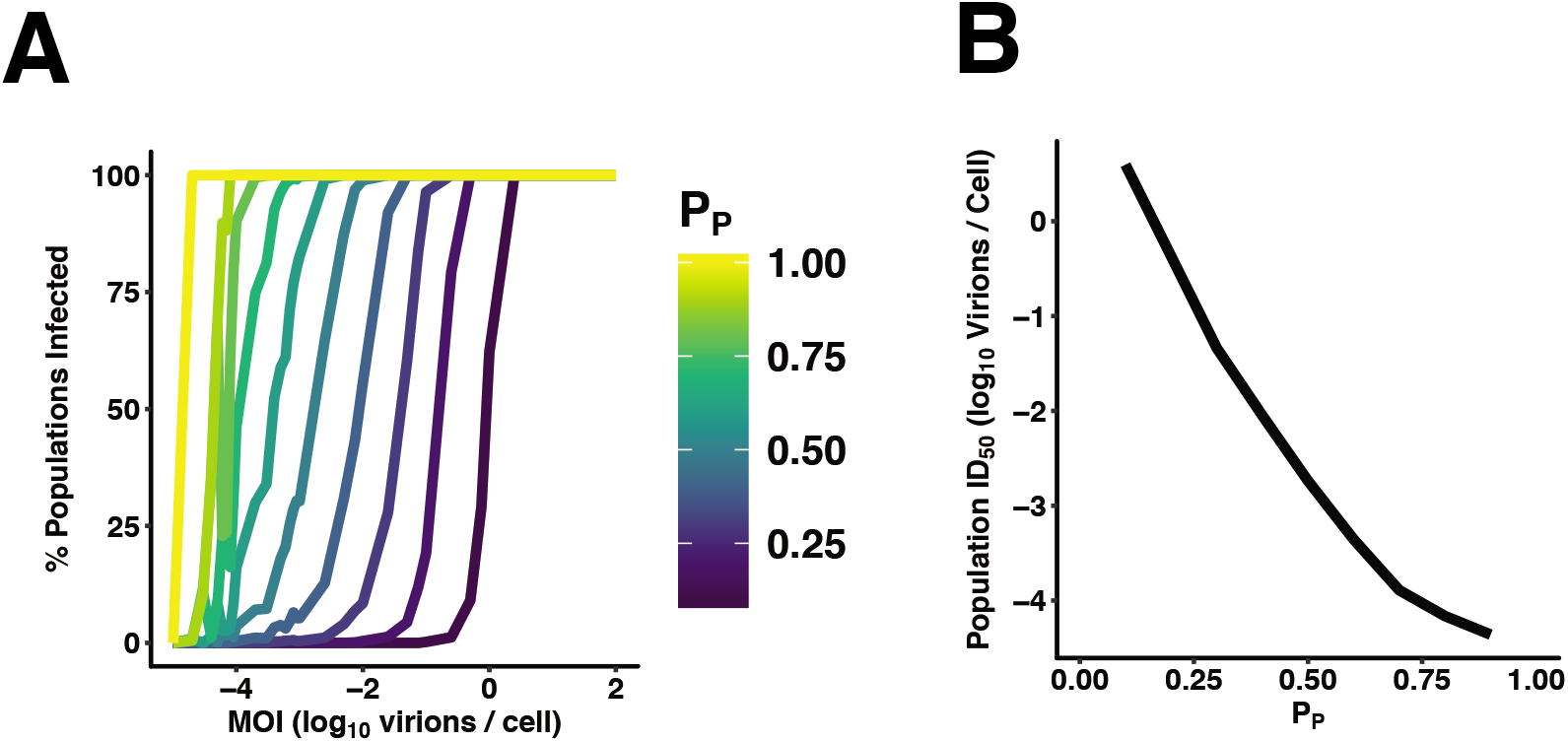
Requirement for co-infection poses a barrier to establishing an infection in a population of cells. To define the impact of IVGs on the ability of a virus population to establish infection in a naïve population of cells, simulations were conducted in which varying numbers of virus particles with varying P_P_ values infected a monolayer of cells (N = 100 simulations per (P_P_, MOI) combination). (A) At each PP and MOI, the percentage of populations in which at least one cell contained 8 segments following the inoculation was calculated. (B) The MOI that led to 50% of cell populations becoming infected (ID_50_) was plotted as a function of P_P_.

### Model of spatially structured viral spread

The estimates of viral infectivity made above assume that virus is distributed randomly over a population of cells. Following the initial infection event, however, viruses spread with spatial structure. We hypothesized that this structure may be very important for reducing the costs of genome incompleteness once infection is established. To test this idea, we developed a model of viral spread in which the extent of spatial structure could be varied.

The system comprises a spatially explicit grid of cells that can become infected with virus. The number and type of segments delivered upon infection is dictated by the parameter P_P_ and, if all eight segments are present, a cell produces virus particles. These particles can then diffuse in a random direction, with the distance traveled governed by the diffusion coefficient (D). *D* was varied in the model to modulate the spatial structure of viral spread: higher *D* corresponds to greater dispersal of virus and therefore lower spatial structure. We simulated replication of two virus strains under a range of diffusion coefficients, one with a frequency of IVGs characteristic of Pan/99 virus (P_P_ = 0.58) and one with complete genomes (P_P_ = 1.0).

Our results point to an important role for spatial structure in determining the efficiency of infection. When P_p_ = 1.0, replication proceeds faster at higher values of *D*, because virus particles reach permissive cells more efficiently (Fig. 4A and Supplementary Fig. 2A). In contrast, when P_P_ = 0.58, replication proceeds fastest at intermediate values of *D* (Fig. 4A and Supplementary Fig. 2B). An intermediate level of spatial structure is optimal for a virus with incomplete genomes for two reasons. At high values of *D*, virions diffuse farther and cellular co-infection becomes less likely, reducing the likelihood of complementation. At the other extreme, when *D* is very low, complementation occurs readily but spread to new cells becomes rare.

**Figure 4.**
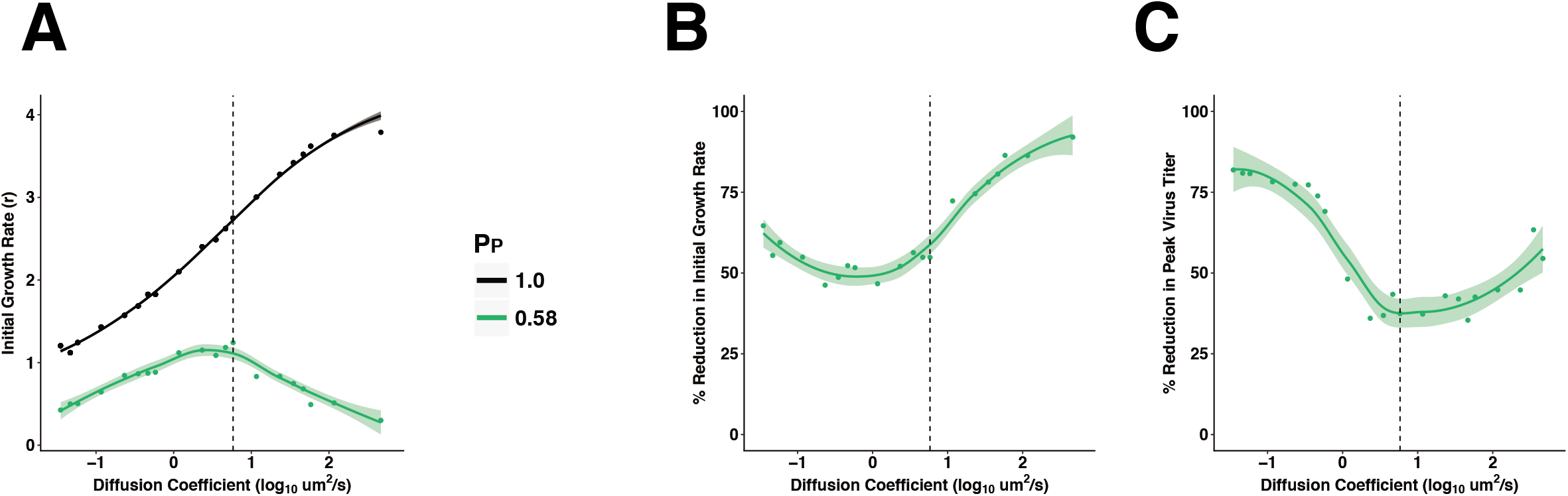
The fitness costs of incomplete genomes may be mitigated by spatially structured spread. The dynamics of multi-cycle replication in a 100×100 grid of cells were simulated, starting from a single cell in the center of the grid. (A) The initial growth rate (estimated by the number of cells infected in the first 12 h) is shown across a range of diffusion coefficients for a virus with P_P_ = 1.0 (black) and P_P_ = 0.58 (green). (B, C) The fitness cost of IVGs, as measured by the reduction in initial growth rate (B) and the reduction in the number of virions produced at peak (C) are shown across a range of diffusion coefficients. The vertical dashed line represents the estimated value of *D* (5.825 um^2^/s) for a spherical IAV particle in water. Each point shows the mean of 10 simulations. Curves were generated by local regression. Shading represents 95% CI.

The model allows the potential costs of incomplete genomes to be evaluated by comparing results obtained for a virus with P_P_=1.0 to those obtained for a virus with a lower P_P_. In particular, we focused on P_P_=0.58 based on the measured values for Pan/99 virus. In Figure 4B and 4C we show how reductions in the initial growth rate (Fig. 4B) and in the amount of virus produced at the peak of infection (Fig. 4C) brought about by incomplete genomes vary with spatial structure. We see that costs of incomplete genomes are minimized at intermediate values of D. These results predict that the fitness effects of IVGs are dependent on the extent to which viral dispersal is spatially constrained.

### Impact of MOI on efficiency of virus production

Burst size, the average number of virions generated by an infected cell, is an important factor determining the potential for complementation of incomplete genomes. If an infected cell produces a larger number of viral progeny, the likelihood of coinfection in neighboring cells increases. We therefore measured this parameter experimentally for Pan/99 virus by performing single cycle growth assays over a range of MOIs (1, 3, 6, 10, and 20 PFU/cell). Multiple MOIs were used to determine whether burst size is dependent on the number of viral genome copies per cell. We saw that higher MOIs resulted in earlier emergence of virus, suggesting that there is a kinetic benefit of additional vRNA input beyond what is required to productively infect a cell (Fig.5A; Supplementary Figure 3). Despite these kinetic benefits, MOIs above 3 PFU/cell conferred no benefit in terms of percent infection (Fig. 5B) or total productivity (Fig. 5C). This growth analysis indicated that a maximum of 11.5 PFU per cell was produced during Pan/99 virus infection of MDCK cells. Based on measured P_P_ values, these data estimate that a single productively infected cell produces 962 virions, and this value was used as the burst size in our models.

**Figure 5.**
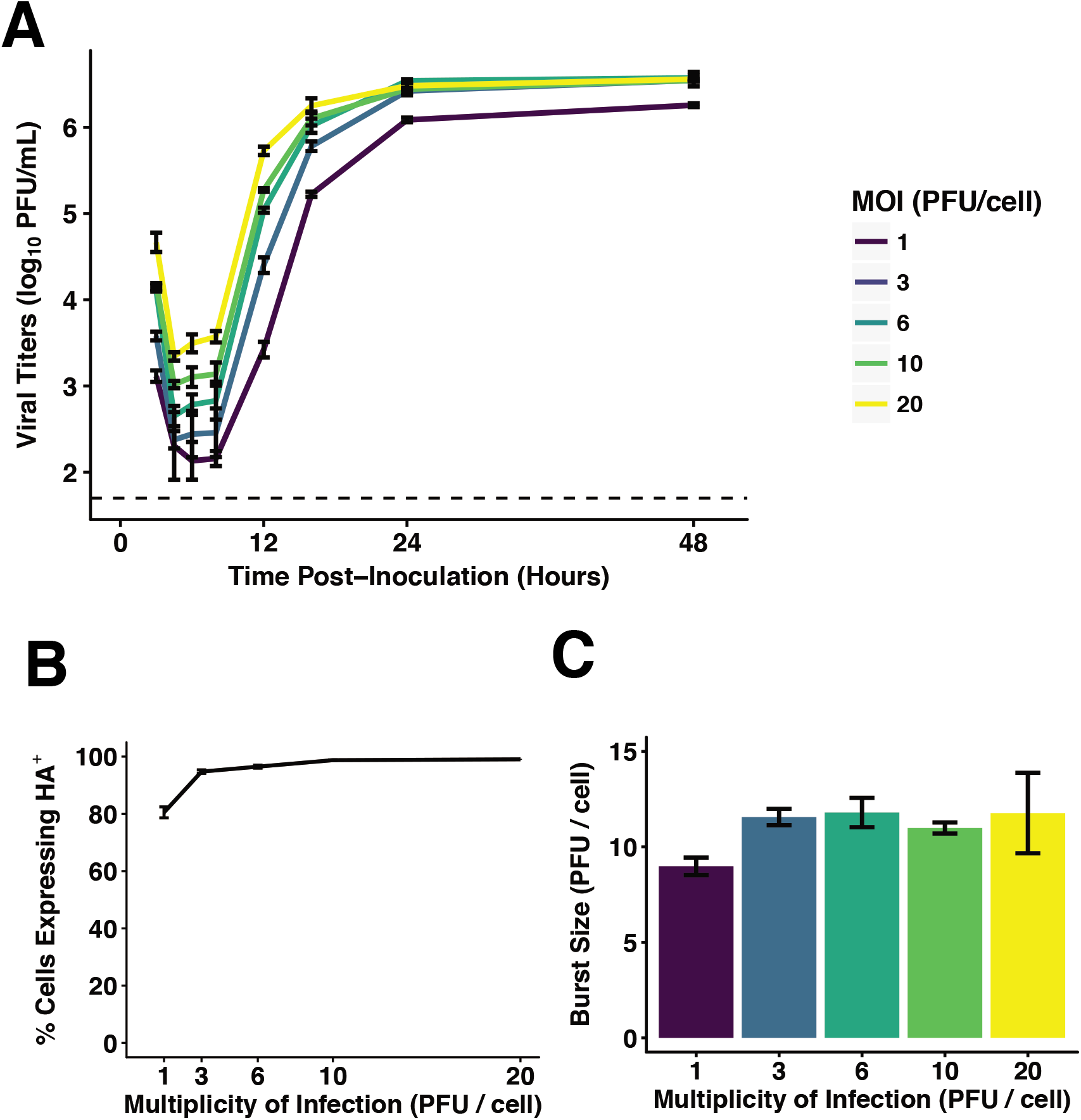
Burst size of Pan/99 virus is constant over a range of high MOIs. (A) MDCK cells were inoculated with Pan/99-WT virus at MOIs of 1, 3, 6, 10, and 20 PFU/cell under single-cycle conditions. Infectious titers at each time point are shown, with MOI indicated by the colors defined in the legend. Dashed line indicates the limit of detection (50 PFU/mL). (B) Fraction of cells expressing HA, as measured by flow cytometry, at each MOI. (C) Burst size in PFU produced per HA^+^ cell. In all panels, mean and standard error are plotted and colors correspond to the legend in panel A.

### Impact of MOI and spatial structure on IVG complementation in cell culture

Our models indicate that, for a virus of a given P_P_, the frequency of infected cells containing IVGs is reduced i) at higher MOIs and ii) under conditions of high spatial structure. These predictions can be seen in Figure 6, where we examined how the proportion of infected cells that are semi-infected varies with MOI (Fig. 6A) and with the diffusion coefficient (Fig. 6B). We tested these predictions of the models experimentally by modulating MOI and spatial spread in IAV infected cell cultures and gauging the impact of each manipulation on levels of IVGs.

**Figure 6.**
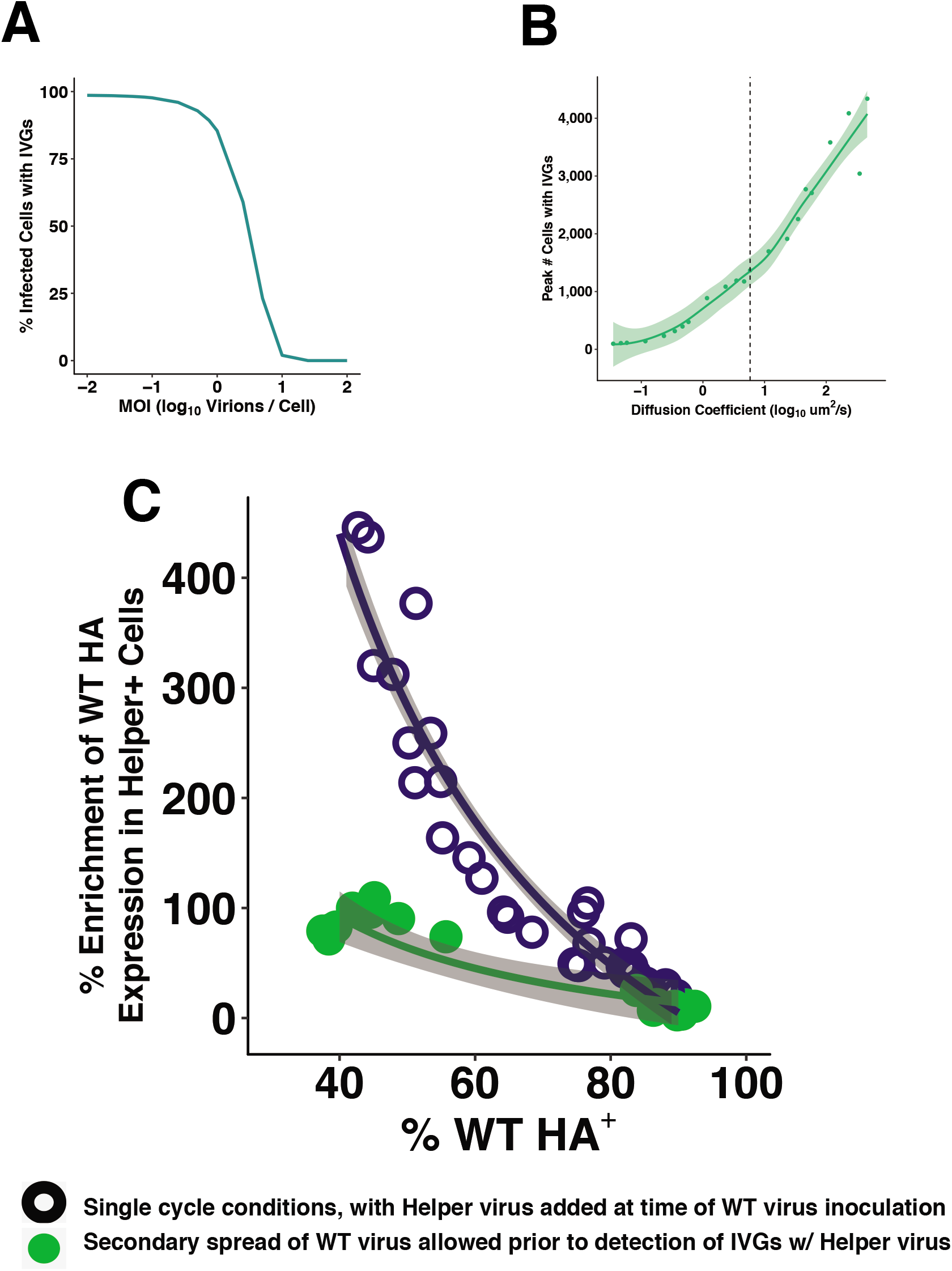
Complementation of incomplete genomes occurs efficiently at high MOI and during secondary spread from low MOI. (A) A well-mixed inoculation was simulated as in Fig. 3. The percentage of infected cells that contain fewer than 8 segments is shown at a range of MOIs for P_P_ = 0.58. (B) An infection in which multi-cycle replication occurs with spatial structure was simulated as in Fig. 4. The maximum number of cells that become semi-infected in each simulation is shown for a range of diffusion coefficients. (C) The extent to which the presence of Pan/99-Helper virus increased WT HA positivity (% Enrichment) was evaluated at the outset of infection (open circles) and following secondary spread (filled circles). To gauge potential for complementation at the outset of infection, cells were simultaneously inoculated with Pan/99-WT virus and Pan/99-Helper, then incubated under single-cycle conditions for 12 h. To test the impact of secondary spread on potential for complementation, cells were inoculated with Pan/99-WT at low MOI and incubated under multi-cycle conditions for 12 h, then inoculated with Pan/99-Helper and incubated under single-cycle conditions for 12 h. Shading represents 95% CI. IVGs = incomplete viral genomes

To monitor levels of IVGs, we used flow cytometry to measure the potential for complementation—that is, the benefit provided by the addition of Pan/99-Helper virus. We hypothesized that, under single cycle conditions, the potential for complementation would decrease with increasing WT virus MOI, since complementation between co-infecting WT viruses would occur frequently at high MOIs. In addition, under multicycle conditions initiated from low MOI, we predicted that the potential for complementation would be greatest at the beginning of infection, due to the random distribution of viral particles, and reduced by secondary spread. We hypothesized that the combination of local dispersal and high particle production during secondary spread would support co-infection in neighboring cells. To test our hypotheses, we inoculated cells with Pan/99-WT virus and either added Pan/99-Helper virus at the same time, or added the Helper virus after allowing time for secondary spread.

To assess the potential for complementation at the outset of infection and at a range of MOIs, cells were co-inoculated with Pan/99-Helper virus at a constant MOI and with Pan/99-WT virus at MOIs of 0.1, 0.3, 0.6, or 1 PFU/cell. Cells were then incubated under single-cycle conditions for 12 h to allow time for HA protein expression. Samples were processed by flow cytometry with staining for WT and Helper HA proteins (Supplementary Figure 4). In each co-infection, we quantified the benefit provided by Pan/99-Helper virus by calculating the enrichment of WT HA expression in Helper^+^ cells relative to Helper^−^ cells. Essentially, the enrichment measure works as follows. If the proportion of Helper^+^ cells that are WT^+^ is higher than the proportion of Helper^−^ cells that are WT^+^, enrichment will be > 0%, indicating a cooperative interaction in which Helper virus allows the expression WT HA genes present in semi-infected cells. The results shown in Figure 6C revealed that the potential for complementation at the outset of infection was high at low MOIs, but decreased with increasing MOI. This result was as expected, since complementation between WT virus particles was predicted to reduce the need for Helper virus (Fig. 6A).

To evaluate the impact of spatially structured secondary spread on IVG prevalence, cells were inoculated with Pan/99-WT virus at low MOI (0.002 or 0.01 PFU/cell) and then multicycle replication was allowed to proceed over a 12 h period. After this period, cells were inoculated with Pan/99-Helper virus to complement any semi-infected cells, and incubated for 12 h under single-cycle conditions to allow HA expression to occur. In contrast to the results seen when complementation was offered at the outset of infection, the enrichment of WT^+^ cells in the Helper^+^ fraction was significantly lower in these samples where multi-cycle replication occurred prior to the addition of Helper virus. This reduction in enrichment is clear when comparing infections performed under each condition in which ~50% of cells expressed WT HA (Fig. 6C). These data agree with our theoretical results (Fig. 6B) and indicate that the spatial structure of secondary spread facilitates complementation between WT particles as they infect neighboring cells at locally high MOIs.

### Generation of a virus with absolute dependence on multiple infection

To evaluate the potential for complementation *in vivo*, we generated a virus that is fully dependent on complementation for replication. This was accomplished by modifying the M segment to generate one M segment which encoded only M1 (M1.Only) and a second one which encoded only M2 (M2.Only) (Fig. 7A). When combined with seven standard reverse genetics plasmids for the remaining viral gene segments, the plasmids encoding these two M segments allowed the generation of a virus population in which individual viruses encode functional M1 or M2, but not both. We called this virus Pan/99-M.STOP virus. Due to the rarity of recombination within segments in negative-sense RNA viruses^30^, it is unexpected that M1.Only and M2.Only segments will recombine to generate a WT M segment. Hence, this virus is reliant on both M segments being delivered to the same cell by co-infection. This requirement for co-infection is only absolute *in vivo*, as M2 is not essential for replication in cell culture^31, 32^. However, this virus is expected to be attenuated in cell culture because a lack of M2 hinders replication^33^, and also because some proportion of virions contain M segments that do not encode M1 and so will be unable to replicate independently. It is important to note that, in contrast to the more arbitrary multiplicity dependence of a wild type IAV, the complementation needed by Pan/99-M.STOP virus requires co-infection with two viruses of a particular genotype.

**Figure 7.**
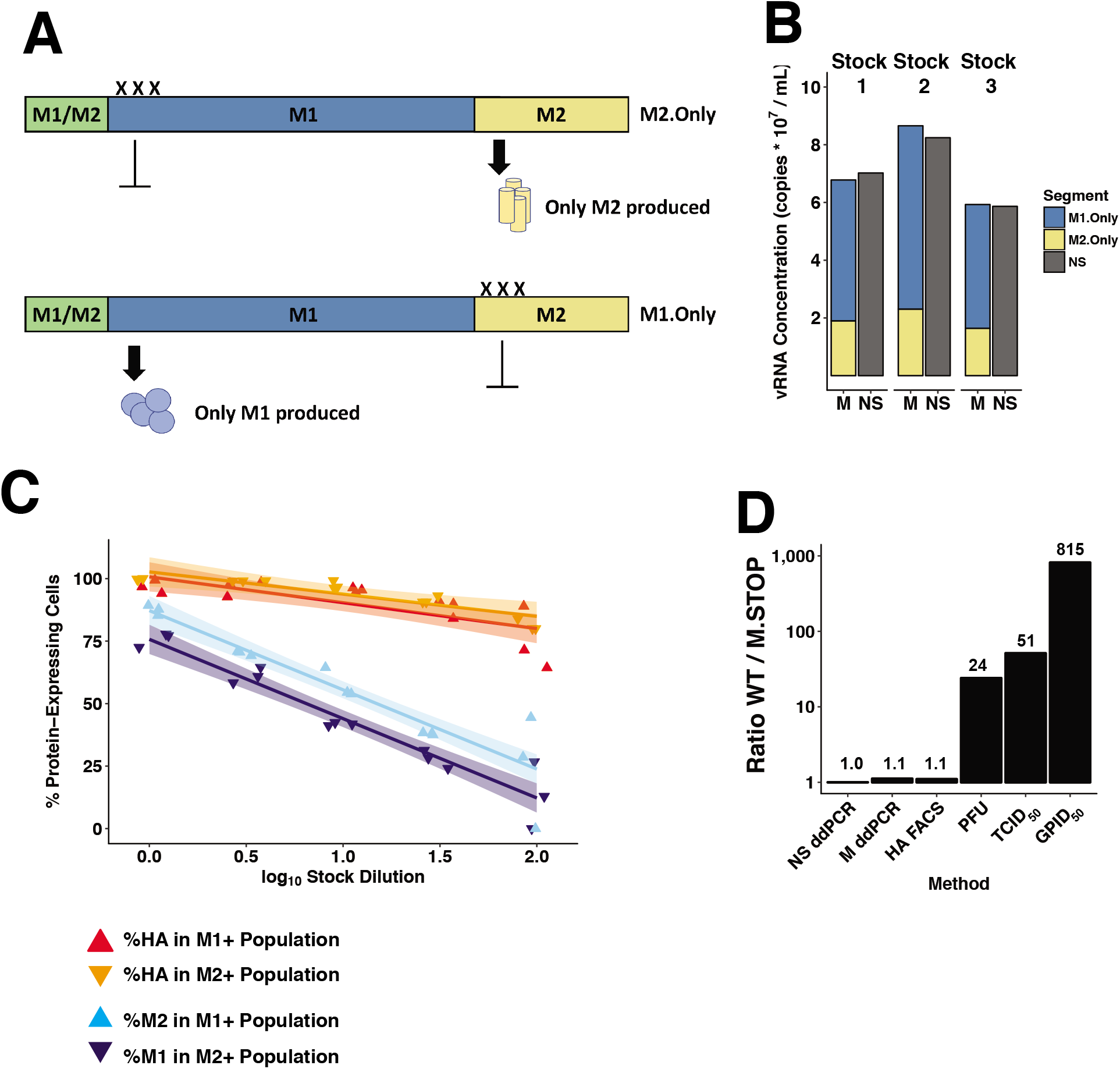
Dependence on complementation hinders viral infectivity. (A) Mutation scheme used to generate M1.Only and M2.Only segments. (B) Copies of M1.Only, M2.Only, and NS segments in three separate Pan/99-M.STOP virus stocks were quantified by digital droplet PCR. (C) Cells were inoculated with Pan/99-M.STOP virus and incubated under single-cycle conditions before staining for HA, M1, and M2 expression. The percentage of cells expressing M1, M2, and HA within M1^+^ or M2^+^ subpopulations is shown at each dilution. Curves represent linear regression with shading representing 95% CI. (D) Titers of WT and M.STOP virus stocks were quantified by ddPCR targeting the NS segment, ddPCR targeting (any) M segment, immunotitration by flow cytometry, plaque assay, tissue culture ID_50_, and guinea pig ID_50_. All results are normalized to the ratio of NS ddPCR copy numbers.

To characterize the Pan/99-M.STOP virus genetically, we used digital droplet PCR (ddPCR) to measure copy numbers of the two M segments and the NS segment in three different virus stocks (Fig. 7B). The total M segment copy number was found to comprise 30% M2.Only and 70% M1.Only. In addition, the total number of M segments was similar to the number of NS segments, as expected if each virion packages one NS and one M vRNA (Fig. 7B). To verify that M1.Only and M2.Only M segments were packaged into distinct virions, we performed infections of MDCK cells with serial dilutions of Pan/99-M.STOP virus under single-cycle conditions and analyzed expression of M1 and M2 by flow cytometry. We observed that, as dilution increased, cells expressing M1 were less likely to express M2, and vice versa (Fig. 7C). This result would be expected if expression of both proteins from the same cell required co-infection with M1.Only and M2.Only encoding virions. As a control, we monitored the effect of dilution on co-expression of HA and M1 or M2. Here, we found that co-expression of M1 or M2 and HA was much less sensitive to dilution, consistent with co-delivery of M and HA segments by single virions.

To test the hypothesis that a given number of Pan/99-M.STOP virus particles would be less infectious than a comparable number of Pan/99-WT virus particles, we characterized both viruses using a series of titration methods that vary in their dependence on infectivity and M protein expression. We first used ddPCR to quantify NS copy numbers of the WT and M.STOP viruses and then normalized all other comparisons to this ratio to account for the difference in virus concentration. As shown above, total M copy numbers were roughly equivalent when normalized to NS (Fig. 7B and 7D). Using immunotitration, in which cells are infected under single-cycle conditions with serial dilutions of virus and then stained for HA expression^34^, we observed equivalent titers of both viruses (Fig. 7D). This was expected, as HA expression under single-cycle conditions is not dependent on M1 or M2 proteins. When titration relied upon multi-cycle replication, however, the WT virus was higher titer than the M.STOP virus. This difference was moderate in cell culture-based measurements, with PFU and TCID_50_ titers 24- and 51-fold higher, respectively, likely because of the reduced importance of M2 in this environment. The full cost for infectivity of separating the M1 and M2 ORFs onto distinct segments was apparent *in vivo*, where 815-fold as much M.STOP virus was required to infect 50% of guinea pigs compared to WT virus (Fig. 7D). Thus, although the M.STOP virus differs from a virus with very low P_P_ in that complementation can only occur when viruses carrying M1.Only and M2.Only segments co-infect, the prediction shown in Figure 3 that increased dependence on multiple infection decreases infectivity held true in this system.

### *Potential for complementation* in vivo

Having determined that the dependence of Pan/99-M.STOP virus on complementation impairs viral infectivity, we next sought to evaluate the potential for complementation to occur *in vivo* once infection had been established. Guinea pigs were inoculated intranasally with equivalent doses of Pan/99-WT or Pan/99-M.STOP virus in terms of NS vRNA copies. Specifically, a dose of 10^7^ copies per guinea pig was used to ensure successful Pan/99-M.STOP virus infection in all animals. This dose represents 8 x GPID_50_ of this mutant virus and 6.5×10^3^ × GPID_50_ of the WT virus. Despite its reduced ability to establish infection, Pan/99-M.STOP virus successfully grew in guinea pigs, following similar kinetics to Pan/99-WT virus. Average peak virus production, measured as NS vRNA copies, was reduced by only 9-fold relative to WT (Fig. 8A).

**Figure 8.**
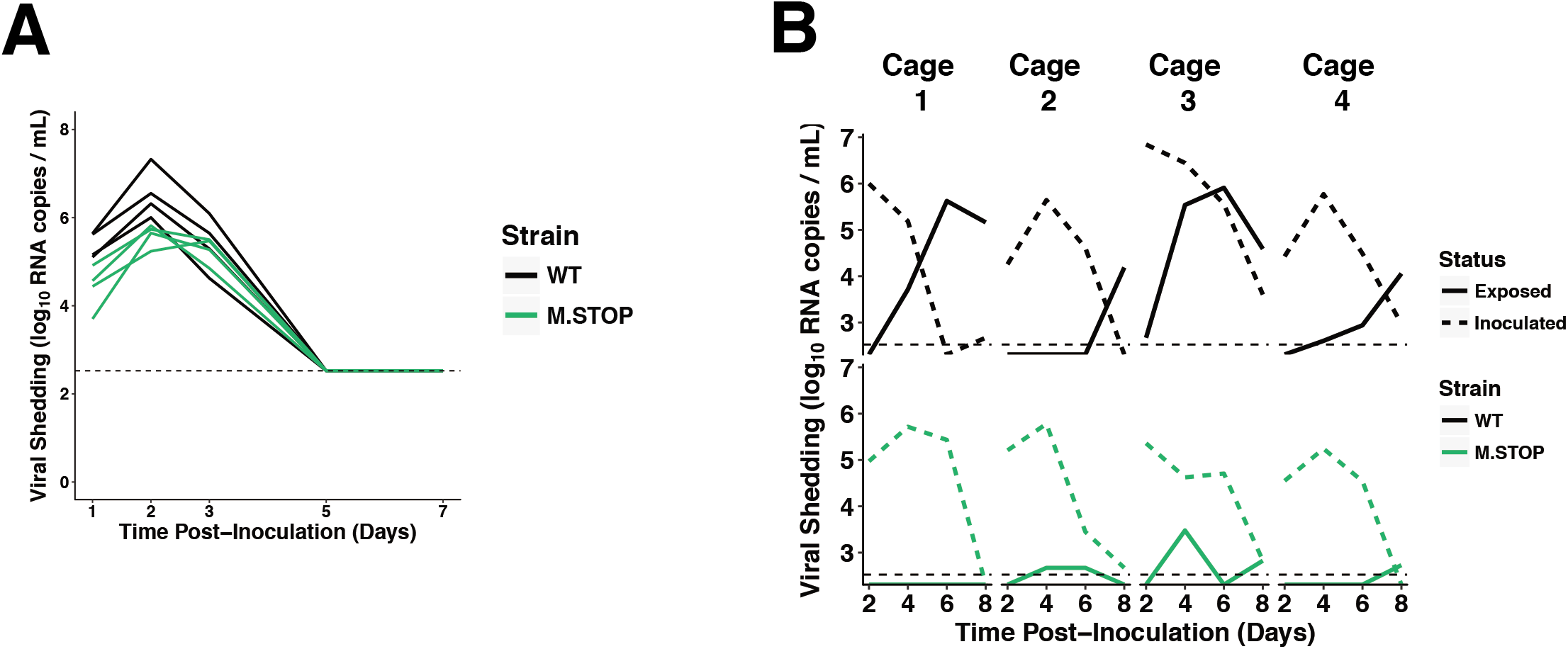
Dependence on complementation hinders viral transmission but has a more modest effect on replication. (A) Guinea pigs were inoculated with 10^7^ RNA copies of Pan/99-WT virus or Pan/99-M.STOP virus, and nasal washes were collected over 7 days to monitor shedding. NS segment copy number per mL of nasal lavage fluid is plotted. (B) Guinea pigs were inoculated with 8 x GPID_50_ of Pan/99-WT virus or Pan/99-M.STOP virus and co-housed with uninfected partners after 24 h. Nasal washes were collected over the course of 8 days to monitor shedding kinetics and transmission between cagemates. NS segment copy number per mL of nasal lavage fluid is plotted. Horizontal dotted line represents the limit of detection (335 RNA copies/mL).

Next, we set up an experiment to determine whether Pan/99-M.STOP virus was competent to undergo transmission to new hosts. In this case, guinea pigs were inoculated with equivalent doses in terms of GPID_50_ with the goal of establishing comparable infections in the donor hosts so that relative efficiency of transmission could be better evaluated. Thus, doses of 8 × GPID_50_ of WT or M.STOP virus were used. At 24 h post-inoculation, each index guinea pig was co-housed with a naïve partner. As expected, WT virus transmitted to and initiated robust infection in of the four contact animals. In contrast, only transient, low levels of the M.STOP virus was observed in nasal washings collected from contacts (Fig. 8B). These results suggest that the spatial structure inherent to multi-cycle replication mitigates the cost of incomplete genomes in an individual host, but dependence on complementation is costly for transmission.

## Discussion

Using a novel single-cell approach that enables robust detection of incomplete IAV genomes, we show that ~99% of Pan/99 virus infections led to replication of fewer than eight segments. The theoretical models we describe predict that the existence of IVGs presents a need for cellular co-infection, and that this need has a high probability of being met when spread occurs in a spatially structured manner. Use of silent genetic tags allowed us to experimentally interrogate cooperation at the cellular level to test these predictions. In agreement with our models, experiments in cell culture showed that co-infection and complementation occur readily when multiple rounds of infection are allowed to proceed with spatial structure. The high potential for complementation to occur *in vivo* was furthermore revealed by the robust within-host spread of a virus that is fully dependent on co-infection. Complementation was not observed during transmission, however, suggesting that fully infectious particles may be required to initiate infection in a new host.

The existence of incomplete genomes was previously predicted by Heldt et al., and these predictions are consistent with the experimental findings of our single-cell assay^20^. The parameter estimated by this assay, P_P_, is defined as the probability that, following infection with a single virion, a given genome segment is successfully replicated. Previous work by Brooke et al. has shown that cells infected at low MOIs express only a subset of viral proteins^19^. While this failure of protein expression could be explained by a failure in transcription or translation, the results of our single-cell sorting assay indicate that the vRNA segments themselves are absent, as they should be amplified by the helper virus polymerase even if they do not encode functional proteins. As in Brooke et al., our method does not discriminate between the alternative possibilities that segments are absent from virions themselves or are lost within the cell, but published results suggest that a single virion usually contains a full genome^22, 23^. Importantly, our single cell assay quantifies the frequencies of all eight segments, rather than only those that can be detected indirectly by staining for protein expression, and therefore allows for analysis of the associations between segments. Despite the importance of interactions among vRNP segments during virion assembly^10, 26–29^, we did not detect compelling evidence of segment co-occurrence at the level of vRNA replication within target cells. This observation suggests that interactions among segments formed during assembly are likely not maintained throughout the early stages the viral life cycle.

The results of our single cell assay indicate that 1.3% of Pan/99 virions are fully infectious, which is consistent with our prior estimates based on observed levels of reassortment between Pan/99 wild type and variant viruses^24^. This result is, however, lower than other reported estimates of the frequency of fully infectious particles. This difference is likely due in part to our use of a different virus strain, as Brooke et al. observed that this frequency is strain-specific^19^. In addition, our use of a helper virus likely allows more robust detection of IVGs than would be expected in a system dependent on the detection of non-replicating viral genomes or their mRNA transcripts^20, 35^.

Replication and secondary spread in an individual host involves inherent spatial structure, as virions emerge from an infected cell and travel some distance before infecting a new cell^36–38^. Our theoretical model predicts that local co-infection resulting from this spatial structure mitigates the fitness costs of incomplete genomes, but that there is a trade-off between complementation and dispersal. Handel et al. explore a similar trade-off related to attachment rates in well-mixed (unstructured) populations, and find that an intermediate level of “stickiness” is optimal—virions that bind too tightly are slow to leave the cell that produced them, while those bind too weakly are unable to infect new cells^39^. We observe a similar effect with spatial structure: virions that diffuse faster, and hence disperse farther before infecting a new cell, are less likely to co-infect with enough virions to establish a productive infection. By contrast, when virions diffuse more locally, co-infection occurs more frequently than is required for productive infection and virions take longer to physically reach new cells, ultimately limiting spread. The optimal level of spatial structure for a virus with incomplete genomes is thus an intermediate one that allows a population of virions to efficiently reach new cells while ensuring enough complementation to minimize the frequency of semi-infection.

In quantitative terms, our model predicts that a diffusion coefficient characteristic of a sphere of 100 nM diameter in water would give a near-optimal level of spatial structure. While this condition may approximate conditions for spherical virions in cell culture, the extracellular environment experienced by a virus *in vivo* would be different. Namely, virus replicating within the respiratory tract would be released into a layer of watery periciliary fluid, which underlies a more viscous mucous blanket^40^. The structure and composition of this epithelial lining fluid may act to limit dispersal the of virus particles relative to that expected in cell culture. Importantly, however, this fluid lining the airways is not static, but rather is moved in a directional manner by coordinated ciliary action^40^. This coordinated movement raises the interesting possibility that IAVs may have evolved to depend upon ciliary action to mediate directional dispersal of virions to new target cells, while maintaining a high potential for complementation of IVGs. This concept will be explored in subsequent studies.

Our experiments designed to test the predicted role of spatial structure in enabling complementation confirmed that secondary spread allows Pan/99-WT virus to replicate efficiently even at low initial MOIs, diminishing the need for complementation after only 12 hours of multi-cycle replication. The potential for complementation of IVGs *in vivo* was furthermore evidenced by the replication in guinea pigs of Pan/99-M.STOP virus, which requires co-infection for productive infection. Importantly, however, Pan/99-M.STOP virus did not initiate productive infection in exposed cagemates. In interpreting this result, it is important to note that the complementation needed by Pan/99-M.STOP virus requires co-infection with two viruses of a particular genotype. This type of complementation has a lower probability of occurring than that typically needed for completion of a WT IAV genome. Despite this caveat, the failure of Pan/99 M.STOP virus to transmit suggests that the establishment of IAV infection requires at least some fully infectious virions. The delivery of multiple particles to a small area via droplet transmission may allow multiple virions to infect the same cell and establish infection, but our data suggest that this mechanism does not occur efficiently in a guinea pig model. The tight genetic bottleneck observed in human-human transmission events is furthermore consistent with a model in which infection is commonly initiated by single particles^41^. In prior work, mutations decreasing the frequency of fully infectious particles, but not eliminating them entirely, were observed to increase transmissibility^25^. This enhanced transmission was attributed to modulation of the HA:NA balance, which enhanced growth in the respiratory tract. In contrast, the HA:NA balance of the Pan/99-M.STOP virus evaluated herein is not expected to differ from Pan/99 virus.

In summary, our findings suggest that incomplete genomes are a prominent feature of IAV infection. These semi-infectious particles are less able to initiate infections in cell culture and during transmission to new hosts, when virions are randomly distributed. In contrast, our data show that incomplete genomes actively participate in the within-host dynamics of infection as they are complemented by cellular co-infection, suggesting an important role for spatial structure in viral spread. This frequent co-infection leads to higher gene copy numbers at the cellular level, consequently promoting reassortment and free mixing of genes. Thus, a reliance of IAVs on co-infection may have important implications for viral adaptation to novel environments such as new hosts following cross-species transmission.

## Acknowledgements

We thank Daniel Perez for the pDP2002 plasmid, and Shamika Danzy for technical assistance. This work was funded in part by the NIH/NIAID Centers of Excellence in Influenza Research and Surveillance (CEIRS), contract number HHSN272201400004C (to ACL and JS), and by NIH/NIAID grants R01 AI099000 (to ACL) and U19 AI117891 (to RA, ACL and JS).

## Author Contributions

NTJ contributed to the conception of work, experimental design, data acquisition and analysis, interpretation of data and writing of the manuscript; NOO contributed to data acquisition and analysis; AA contributed to data analysis and interpretation; JS contributed to experimental design, interpretation of data and writing of the manuscript; RA contributed to conception of work, interpretation of data and writing of the manuscript; ACL contributed to conception of work, experimental design, interpretation of data and writing of the manuscript.

## Computational Methods

### Probabilistic model to estimate costs of incomplete genomes for cellular infectivity

To define the impact of incomplete viral genomes on viral infectivity, we considered how the infectious dose varies with P_P_, the probability that an individual genome segment from an infectious virus is successfully delivered and replicated within the infected cell. This model assumes that a single particle can deliver each of the 8 segments that a host cell does not already contain. Furthermore, delivery of each segment is independent, making the action of segment delivery by a single virion a binomial process (p = P_P_, N = # of missing segments).

We model this process as a Markov chain in which a cell can exist in 9 states, containing between 0 and 8 genome segments, and transitions between states are governed by the 9×9 matrix *T*, in which each element is described by the binomial distribution:

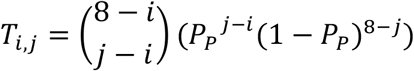

where *i* is the number of segments a cell contains before infection, and *j* is the number of segments it contains after infection. Since the binomial distribution is not defined for *k* < 0, all entries below the main diagonal are populated by 0s. The state of 8 segments, or productive infection, becomes the absorbing state, and it is assumed that each cell will obtain all 8 genome segments given the addition of enough virions. To estimate how many virions are required to reach this state, we first define a 1×9 vector representing the distribution of segments per cell.

To represent an uninfected cell, we set τ_0_ = [1,0…0]. The distribution of segments in a cell that has been infected with *v* virions is then given by:

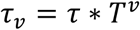

With the element of τ_*v*_ representing the probability a cell contains 8 segments and is therefore productively infected.

Finally, we use survival analysis to calculate the expected number of virions that must infect a cell before it receives all 8 segments:

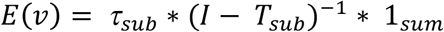

wherein *T_sub_* represents the upper-left 8×8 matrix of *T* (in which a cell contains 0 – 7 segments), and τ_sub_ is the first 8 columns of τ_0_, *I* is the identity matrix, and 1_*sum*_ is an 8×1 vector that acts to sum each state into a single value. This summary statistic represents the number of transitions required for a cell to leave the semi-infected state or, more simply, the average number of virions required to infect a cell, which we define as the “infectious unit.”

To define the impact of semi-infectious particles on the ability of a virus population to establish infection in a naïve population of cells, Monte Carlo simulations were conducted in which varying numbers of virus particles infected a monolayer of cells in a Poisson-distributed manner, with different P_P_ values. 1000 simulations were run per (P_P_, MOI) combination. At each P_P_ and MOI, the percentage of cells containing 8 segments and the percentage of cells containing between 1 and 7 segments were recorded, as well as the percentage of populations in which at least one cell contained 8 segments following the inoculation. The MOI that led to 50% of cell populations becoming infected (ID_50_) was calculated for each P_P_ value based on four-parameter logistic regression of the dose-response curves shown in Figure 3A.

### Individual-based model of replication

A cellular automaton model of viral spread was developed to investigate the relationships among spatial structure, prevalence of incomplete viral genomes, and viral fitness. The system consists of a 100×100 grid of cells. Each cell contains 0 – 8 unique IAV genome segments. Virions exist on the same grid, in a bound or unbound state. When a virion infects a cell, any missing segments may be delivered, with the probability of delivery defined by P_P_, as derived in Figure 1. The simulation begins with a single productively infected cell in the middle of the grid. The following events occur at each time-step (1 minute), and the frequency of each of these events is governed by the parameters listed in Table 1.
1. Productive cells (containing 8 segments) produce virions, which are initially bound to the producer cell’s surface.
2. Free virions may attach to the cell at their current position.
3. Bound virions may infect the cell to which they are attached, or be released.
4. Free virions diffuse some normally distributed distance 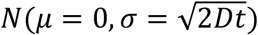, where *D* is the diffusion coefficient, *t* is the length of the time-step, *μ* is the mean and *σ* is the standard deviation.
5. Productive cells may die.
6. Infected cells (1 – 8 segments) may become refractory to super-infection.

To generate the data shown in Figure 4 and Figure 6B, these events were iterated over multiple rounds of infection up to 96 h post-infection. Ten simulations per (*D*, P_P_) combination were conducted.

## Experimental Methods

### Cells

Madin-Darby canine kidney (MDCK) cells (contributed by Peter Palese, Icahn School of Medicine at Mount Sinai) were cultured in minimal essential medium (MEM) supplemented with 10% fetal bovine calf serum (FCS), penicillin (100 IU), and streptomycin (100 ug/mL). 293T cells (ATCC, CRL-3216) were cultured in Dulbecco’s modified essential medium (DMEM) supplemented with 10% FCS.

As used herein, “MEM” refers to MEM supplemented with 10% FCS and penicillin/streptomycin at the above concentrations, which was used for maintaining cells in culture. Following infection with influenza viruses, cells were incubated with “virus medium”, which herein refers to MEM supplemented with 0.3% bovine serum albumin and penicillin/streptomycin at said concentrations. When their presence is indicated, TPCK-treated trypsin was used at 1 ug/mL, NH_4_Cl at 20 mM, HEPES at 50 mM.

### Viruses

All viruses were generated by reverse genetics following modification of the influenza A/Panama/2007/99 (H3N2) virus cDNA, which was cloned into pDP2002 (Chen et al. 2012). All viruses were cultured in 9-11 day old embryonated hens’ eggs unless otherwise noted below. To limit propagation of defective interfering viral genomes, virus stocks were generated either from a plaque isolate or directly from 293T cells transfected with reverse genetics plasmids. The only genetic modification made to the Pan/99-WT virus was the addition of sequence encoding a 6-His tag plus GGGGS linker following the signal peptide of the HA protein as previously described^8^.

A genetically distinct but phenotypically similar virus, referred to herein as “Pan/99-Helper”, was generated by the introduction of six or seven silent mutations on each segment, as well as the addition of the HA-tag (sequence: YPYDVPDYA) instead of the 6-His tag. The silent mutations are listed in Supplementary Table 1 and were designed to introduce strain-specific primer binding sites, allowing the presence or absence of each segment to be measured by qRT-PCR. Epitope tags in HA allowed identification of infected cells by flow cytometry.

A virus with two distinct forms of the M segment, referred to herein as “Pan/99-M.STOP virus”, was constructed. Site-directed mutagenesis was used to introduce nonsense mutations into the pDP2002 plasmid containing the sequence of the M segment in order to abrogate expression of M2 but not M1 (M1.Only), or vice versa (M2.Only). For M2.Only, three in-frame stop codons were introduced downstream of the sequence encoding the shared M1/M2 N-terminus. An in-frame ATG at nucleotide 152 was also disrupted. For M1.Only, three in-frame stop codons were introduced in the M2 coding region downstream of the M1 ORF. In addition, four amino acid changes were made to M2 coding sequence in the region following the splice acceptor site and upstream of the introduced stop codons. Both plasmids were used in conjunction with pDP plasmids encoding the other seven segments to generate a mixed virus population in which each virion contained an M1.Only or M2.Only segment. At 24 h post transfection, 293T cells were washed with 1 mL PBS, then overlaid with 1×10^6^ MDCK cells in virus medium plus TPCK-treated trypsin, and incubated at 33°C for 48 h. Supernatant was used to inoculate a plaque assay, and after 48 h a plaque isolate was used to inoculate a 75 cm^2^ flask of MDCK cells. Following 48 h of growth, this stock was aliquoted and used to inoculate a plaque assay. One plaque isolate was diluted and used to inoculate 10-day-old embryonated chickens’ eggs for a third passage. Experiments were conducted with this egg passage stock.

### Infections

6-well dishes (Corning) were seeded with 4 × 10^5^ MDCK cells in 2 mL MEM, then incubated for 24 h. Prior to inoculation, MEM was removed and cells were washed twice with 1 mL PBS per wash. Inocula containing virus in 200 uL PBS were added to cells, which were incubated on ice (to permit attachment but not viral entry) for 45 minutes. After inoculation, the monolayer was washed with PBS remove unbound virus before 2 mL virus medium was added and plates were incubated at 33°C. For multi-cycle replication, TPCK-treated trypsin was added to virus medium to a final concentration of 1 ug/mL. When single-cycle conditions were required, virus medium was removed after 3 h and replaced with 2 mL virus medium containing NH_4_Cl and HEPES.

### Flow cytometry

At 12 h post-inoculation, virus medium was aspirated from infected cells, and monolayers were washed with PBS. The monolayer was disrupted using 0.05% trypsin + 0.53 mM EDTA in Hank’s Balanced Salt Solution (HBSS). After 15 minutes at 37°C, plates were washed with 1 mL FACS buffer (PBS + 1% FCS + 5 mM EDTA) to collect cells and transfer them to 1.7 mL tubes. Cells were spun at 2,500 rpm for 5 minutes, then resuspended in 200 uL FACS buffer and transferred to 96-well V-bottom plates (Corning). The plate was spun at 2,500 rpm and supernatant discarded. Cells were resuspended in 50 uL FACS buffer containing antibodies at the following concentrations, then incubated at 4° C for 30 minutes:

1. His Tag-Alexa 647 (5 ug/mL) (Qiagen, catalog no. 35370)
2. HA Tag-FITC (7 ug/mL) (Sigma, clone HA-7)

After staining, cells were washed by three times by centrifugation and resuspension in FACS buffer. After the final wash, cells were resuspended in 200 uL FACS buffer containing 7-AAD (12.5 ug/mL) and analyzed by flow cytometry using a BD Fortessa.

This approach was modified slightly when staining for M1 and M2. After staining for His and HA (where indicated), cells were washed once with 200 uL FACS buffer, then resuspended in 100 uL BD Cytofix/Cytoperm buffer and incubated at 4°C for 20 minutes. BD Cytoperm/Cytowash (perm/wash) buffer was added to each well, and cells were spun at 2,500 rpm for 5 minutes. After a second wash, cells were resuspended in 50 uL perm/wash buffer containing antibodies at the following concentrations:

1. Anti-M1 GA2B conjugated to Pacific Blue (4 ug/mL) (ThermoFisher)
2. Anti-M2 14C2 conjugated to PE (4 ug/mL) (Santa Cruz)

Following another 30 minutes of staining at 4° C, cells were washed three times (as described above) with perm/wash buffer, then resuspended in FACS buffer without 7-AAD just prior to analysis on the BD Fortessa.

### Quantification of P_P_ values

A single cell sorting assay was used to measure the frequency with which individual genome segments are delivered to an infected cell. 4*10^5^ MDCK cells were seeded into a 6-well dish, then counted the next day just before inoculation. Cells were then washed 3x with PBS and co-inoculated with the virus of interest (Pan/99-WT, MOI = 0.5 PFU/cell) and helper virus (Pan/99-Helper, MOI = 3.0 PFU/cell) in a volume of 200 uL. Cells were incubated at 33°C for 60 minutes, after which they were washed 3x with PBS, and 2 mL of virus medium was added. After incubation at 33°C for 60 minutes, medium was removed and cells were washed 3x with PBS before addition of Cell Dissociation Buffer (Corning) containing 0.1% EDTA (w/v) to release cells from the plate surface. Cells were harvested by resuspension in MEM, followed by a series of three washes in 2 mL FACS buffer (2% FCS in PBS). Cells were resuspended in PBS containing 1% FCS, 10 mM HEPES, and 0.1% EDTA and filtered immediately prior to sorting on a BD Aria II. After gating to exclude debris and doublets, one event was sorted into each well of a 96-well plate containing MDCK cell monolayers at 30% confluency in 50 uL virus medium containing TPCK-treated trypsin. After sorting, an additional 50 uL of medium was added to a final volume of 100 uL per well, and plates were spun at 1,800 rpm for 2 minutes to help each sorted cell attach to the plate surface. Plates were incubated at 33°C for 48 h to allow outgrowth of virus from this single infected cell.

RNA was extracted from infected cells using a ZR-96 Viral RNA Kit (Zymo Research) as per manufacturer instructions. Extracted RNA was converted to cDNA using universal influenza primers (given in Supplementary Table 2), Maxima RT (Thermo Scientific, 100 U/sample) and RiboLock RNase inhibitor (Thermo Scientific, 28 U/sample) according to manufacturer instructions. After conversion, cDNA was diluted 1:4 with nuclease-free water and used as template (4 uL/reaction) for segment-specific qPCR using SsoFast EvaGreen Supermix (Bio-Rad) in 10 uL reactions. Primers for each segment of Pan/99-WT virus, as well as the PB2 and PB1 segments of Pan/99-Helper virus, are given in Supplementary Table 2, and were used at final concentrations of 200 nM each.

P_P_ was calculated using the following correction factor to account for the frequency of coinfection:

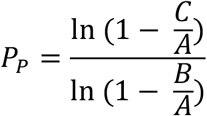

where A is the number of Helper^+^ cells, B is the number of wells positive for any WT segment, and C is the number of wells positive for the WT segment in question.

### Single-cycle growth curves

Cells were inoculated with Pan/99-WT virus at MOIs of 1, 3, 6, 10, or 20 PFU/cell, and incubated with 2 mL virus medium at 33° C. After 3 h, virus medium was replaced with virus medium containing NH_4_Cl and HEPES. 100 uL of medium was collected at 3, 4.5, 6, 8, 12, 18, 24, and 48 h post-inoculation (with replacement by fresh medium to keep volumes consistent) for virus quantification by plaque assay. At 12 h post-inoculation, cells were harvested and stained for analysis of HA expression by flow cytometry.

### Impact of secondary spread on complementation of incomplete genomes

To optimize the approach of using Pan/99-Helper to activate and thereby detect semiinfected cells, we co-inoculated cells with a low MOI (0.01 PFU/cell) of Pan/99-WT virus and a range of Pan/99-Helper virus MOIs and measured expression of WT HA after 12 h. We observed a biphasic relationship between helper virus MOI and the benefit provided to WT virus (Supplementary Figure 3C). As more Helper virus was added, the percentage of cells expressing WT HA initially increased as more cells became co-infected and thus capable of expressing the WT HA protein. But, as the Helper MOI increased further, a competitive effect was observed and the probability of detecting WT HA expression was decreased. Observing that Pan/99-Helper virus provided the greatest benefit—a 2-fold increase in the frequency of WT HA expression—at an MOI of 0.3 PFU/cell, we used that amount in further complementation experiments. Based on measured P_P_ values, this dose is estimated to contain an average of 27 particles/cell.

In single cycle replication conditions, cells were inoculated on ice with Pan/99-WT virus over a range of MOIs (0.1, 0.3, 0.6, 1 PFU/cell) and, at the same time, with Pan/99-Helper virus (MOI = 0.3 PFU/cell) or PBS. After inoculation, cells were washed with 1 mL PBS, 2 mL virus medium (no trypsin) was added, and cells were incubated at 33°C for 3 h, after which initial virus medium was replaced with virus medium containing NH_4_Cl and HEPES. At 12 h post-inoculation, cells were collected and stained for WT and Helper HA expression as described above.

In multi-cycle replication conditions, cells were inoculated on ice with Pan/99-WT at an MOI of 0.01 or 0.002 PFU/cell, and then incubated at 33°C with virus medium containing TPCK-treated trypsin to allow for multi-cycle growth. After 12 h, cells were washed with 1 mL PBS, then inoculated on ice with Pan/99-Helper virus (MOI = 0.3 PFU/cell), or PBS. After inoculation, cells were washed with 1 mL PBS, 2 mL virus medium (no trypsin) was added, and cells were incubated at 33°C for 3 h, after which initial virus medium was replaced with virus medium containing NH_4_Cl and HEPES. At 12 h post-inoculation with Pan/99-Helper virus, cells were collected and stained for WT and Helper HA expression as described above. The amount of complementation provided by Pan/99-Helper virus was calculated using the equation:

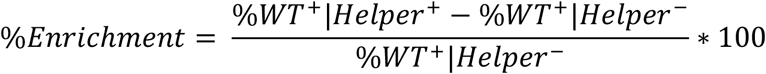

### Digital droplet PCR (ddPCR)

Primers and probes (listed in Supplementary Table 3) were diluted to concentrations of 900 nM and 250 nM per primer and probe, respectively. 22 uL reactions were prepared with 11 uL Bio-Rad SuperMix for Probes (1X final concentration), 6.6 uL of diluted primers (900 nM/primer, final concentration) and probes (250 nM/probe, final concentration), and 4.4 uL of diluted cDNA. 20 uL of each reaction mixture was partitioned into droplets using a Bio-Rad QX200 droplet generator per manufacturer instructions. PCR conditions were: 1.) 95° C for 10 minutes, 2.) 40 cycles of A.) 94° C for 30 seconds and B.) 57° C for 1 minute, 3.) 98° C for 10 minutes, and hold at 4°C. Droplets were then read on Bio-Rad QX200 droplet reader, and the number of cDNA copies/uL was calculated.

### Guinea pig infections

Female Hartley guinea pigs were obtained from Charles River Laboratories (Wilmington, MA) and housed by Emory University Department of Animal Resources. All experiments were conducted in accordance with an approved Institutional Animal Care and Use Committee protocol. For ID_50_ estimation and analysis of viral shedding, guinea pigs were anesthetized by intramuscular injection with 30 mg/kg / 4 mg/kg ketamine/xylazine, then inoculated intranasally with 300 uL virus diluted in PBS. Nasal washes were collected in PBS on days 1, 2, 3, 5, and 7 as described in^42^, and titered by RT ddPCR targeting the NS segment. For transmission experiments, inoculated guinea pigs were individually housed in Caron 6040 environmental chambers at 10°C and 20% relative humidity. At 24 h post-inoculation, one naïve guinea pig was introduced to each cage with one inoculated animal. Nasal washes were collected on days 2, 4, 6, and 8, and titered by plaque assay.

---

**Supplementary Figure 1.**
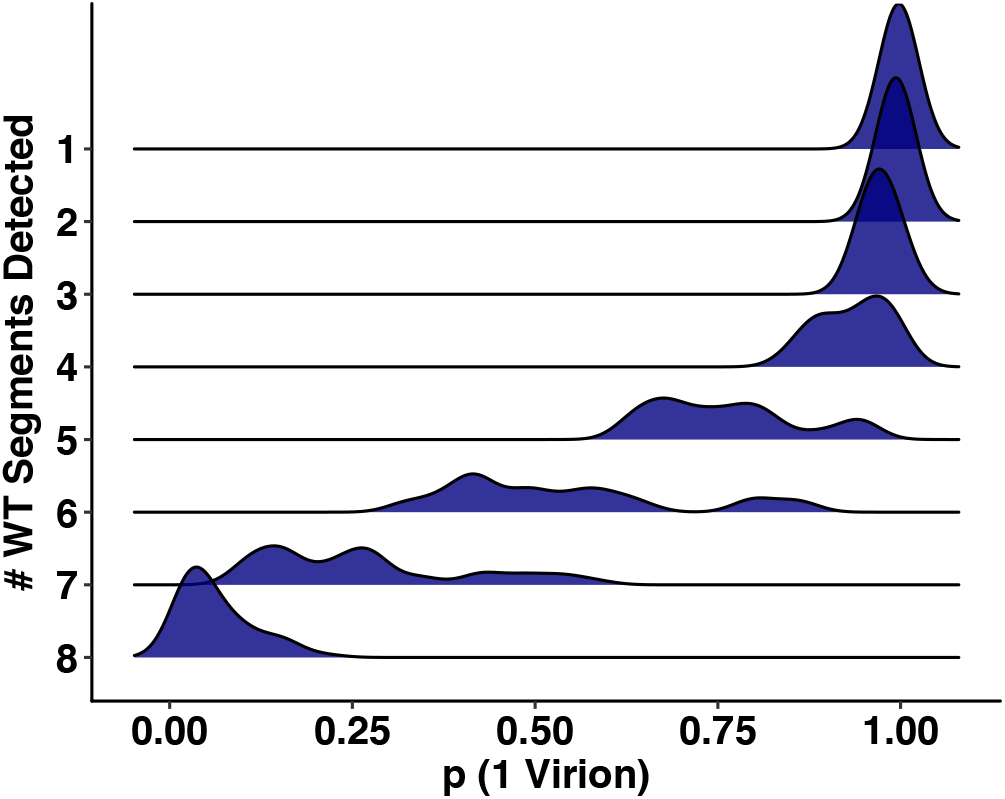
Cells containing more segments were likely to have been infected with multiple virions. Bayes’ rule was used to calculate the probability that each cell was infected with exactly 1 virion, based on the number of infected cells in each experiment, and each cell’s combination of segment presences and absences. The distribution of probabilities is shown stratified by the number of segments present per cell. This figure relates to Figure 1C.

**Supplementary Figure 2.**
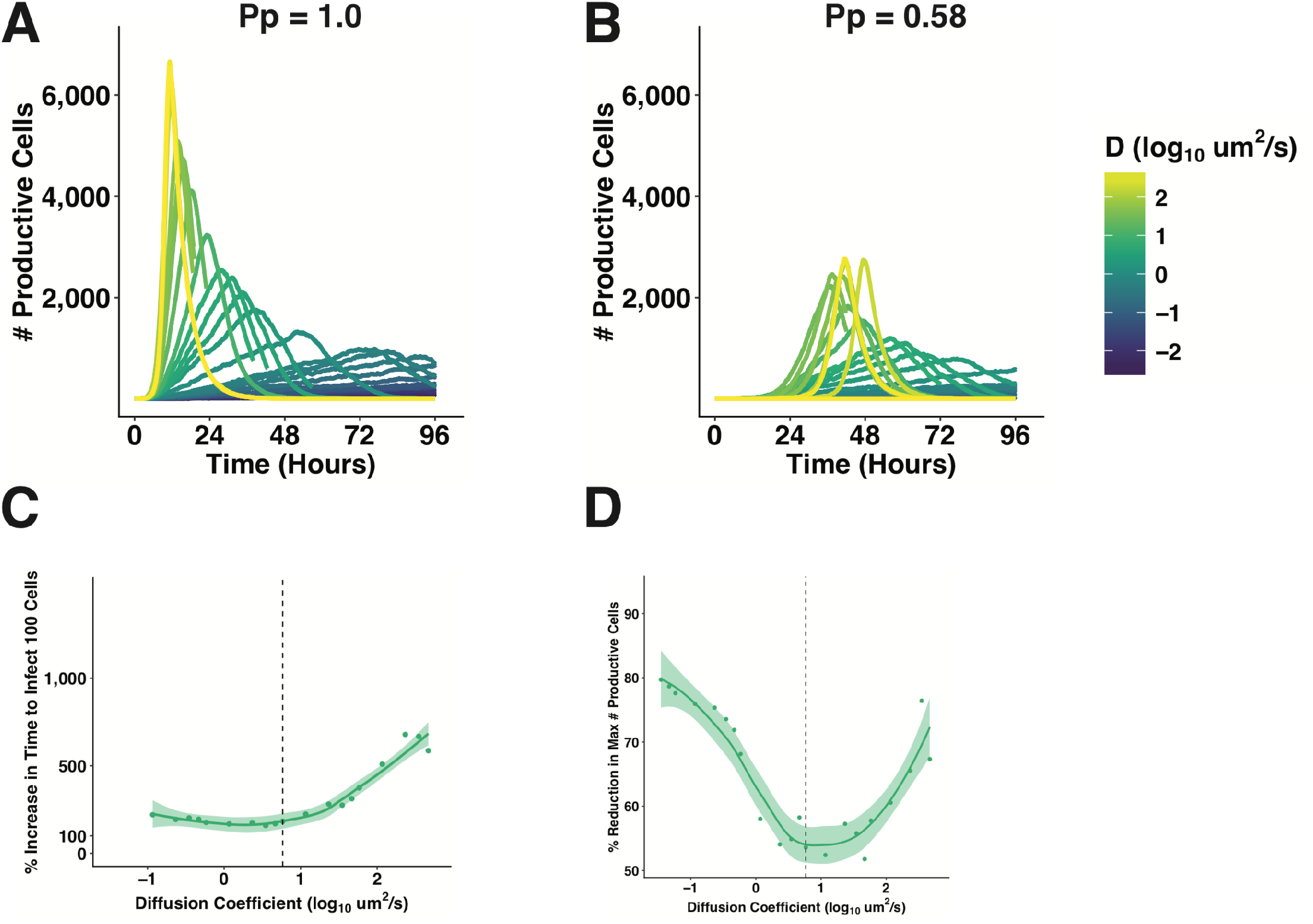
Impact of incomplete genomes and spatial structure on the dynamics and efficiency of IAV infection. (A, B) The dynamics of infection, in terms of productively infected cells, are shown for a virus with no incomplete genomes (P_P_ = 1.0), and a frequency of incomplete genomes similar to Pan/99-WT virus (P_P_ = 0.58). (C, D) The fitness costs of IVGs, in terms of the time taken to productively infect 100 cells (C), and the peak number of virions produced (D), are shown for a range of diffusion coefficients. This figure relates to Figure 4.

**Supplementary Figure 3.**
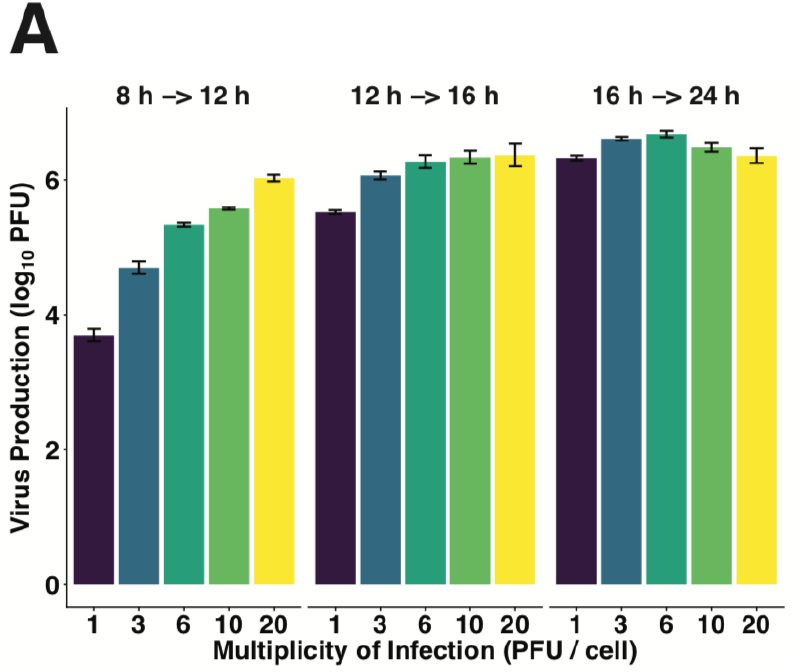
Across a range of high MOIs, the multiplicity of infection impacts the kinetics of viral amplification. The amount of virus produced (in PFU) in three distinct time periods post-infection was calculated at each MOI. This figure relates to Figure 5.

**Supplementary Figure 4.**
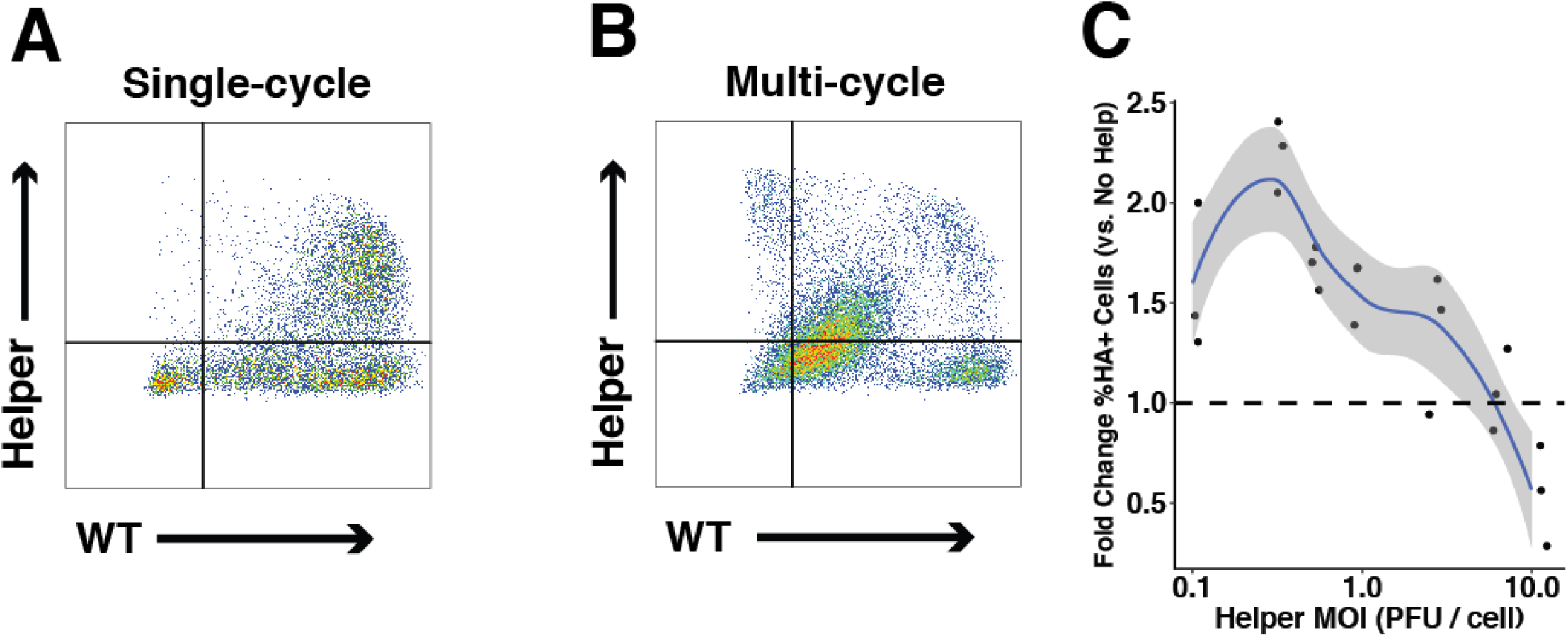
Representative flow plots with staining for WT and Helper HA proteins, and optimization of Helper MOI. (A, B) Representative flow cytometry measurement of Pan/99-WT HA when Pan/99-Helper virus was added simultaneously (A), or following 12 h of multi-cycle replication (B). Pan/99-WT MOI = 0.1 PFU/cell for simultaneous coinfection, 0.01 PFU/cell for multi-cycle replication. Pan/99-Helper virus MOI was 0.3 PFU/cell in both cases. (C) Cells were inoculated with Pan/99-WT (MOI = 0.01 PFU/cell) and Pan/99-Helper at a range of MOI, then incubated under single-cycle conditions before staining for expression of WT and Helper HA proteins. The extent to which Pan/99-Helper increased numbers of WT HA+ cells (relative to controls infected with only Pan/99-WT) was calculated at each Pan/99-Helper MOI. Curve and ribbon represent mean and 95% confidence interval, respectively, of local regression. This figure relates to Figure 6.

**Supplementary Table 1.** Genotype of Pan/99-Helper virus

| **Segment** | **Mutations relative to Pan/99-WT** |
| --- | --- |
| PB2 | A550C, G552A, A555C, C556T, A617C, T621C, T622A, C623G |
| PB1 | C346T, T348G, A351G, T441A, T444A, A447T |
| PA | G603A, T604A, C605G, C747T, T750C, G753A |
| HA | T308C, C311A, C313T, A464T, C467G, T470A |
| NP | C537T, T538A, C539G, G606C, A609T, G615C |
| NA | C418G, T421A, A424C, T511A, T514A, A517G |
| M | C413T, C415G, A418C, A517G, G523A, A526C |
| NS | A210C, G212A, G215A, T218C, C329T, C335T, A341G |

**Supplementary Table 2.**
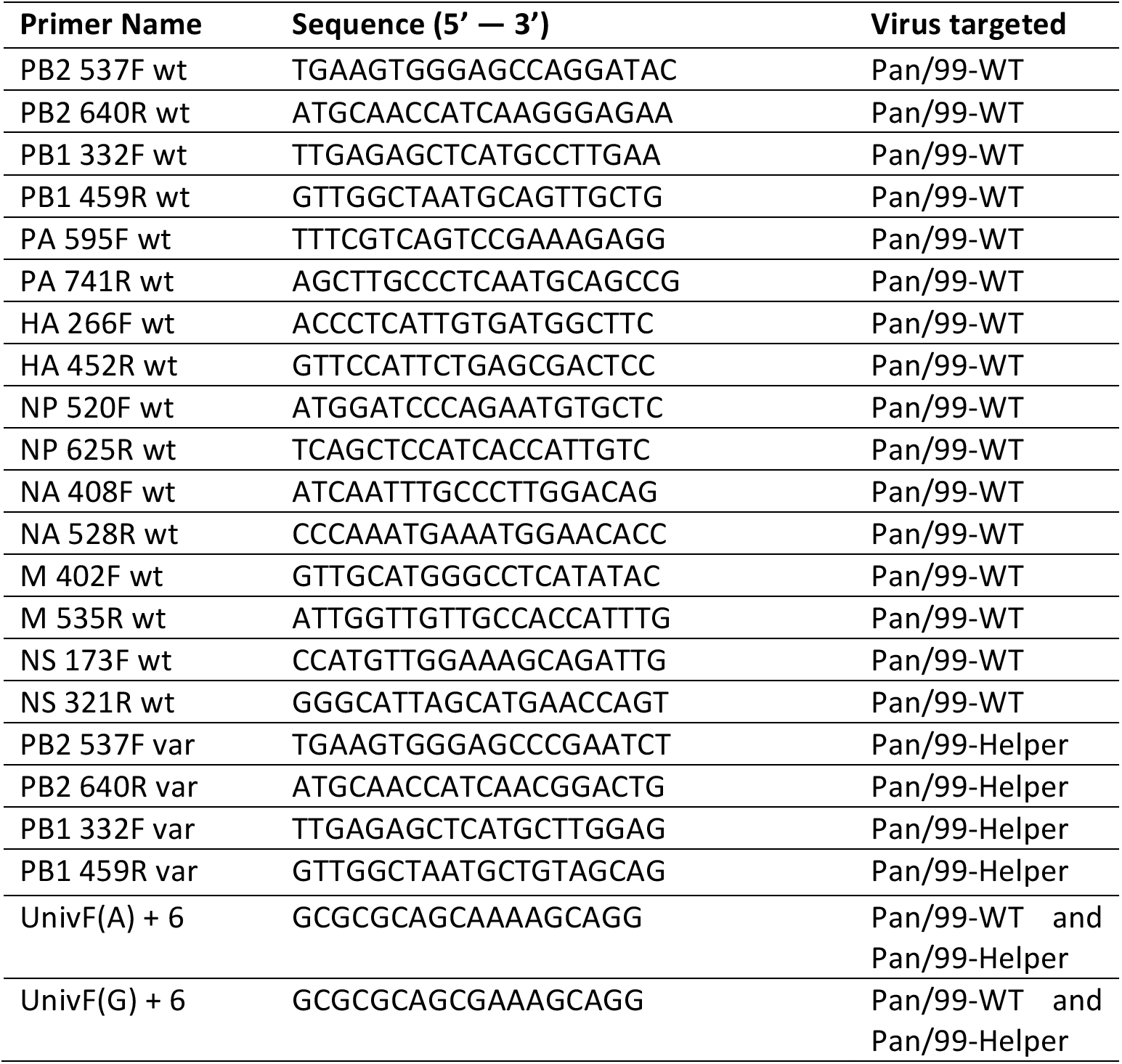
Primers for single-cell assay

**Supplementary Table 3.**
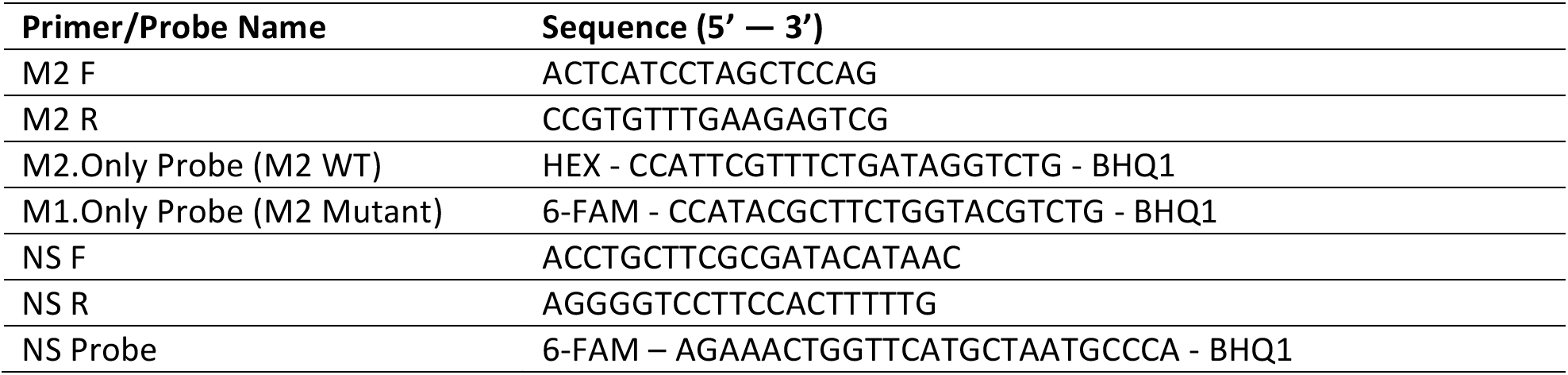
Primers and probes for ddPCR

